# Heat tolerance and its seasonal acclimation in *Fagus sylvatica* compared to *Fagus orientalis* and *Pseudotsuga menziesii*

**DOI:** 10.64898/2026.05.17.725742

**Authors:** Markus Hauck, Germar Csapek, Klara Krämer, Ole Schmidt, Yves Lucas, Lena Popp, Linda Szafranek, Choimaa Dulamsuren

## Abstract

Heat tolerance determines the vitality of tree species under climate change independently of drought tolerance, but has been much less studied than tree water relations. We studied species-specific differences and the capacity for seasonal heat acclimation in Central Europe’s naturally most important tree species, *Fagus sylvatica*, in comparison with two exotic tree species (*Fagus orientalis*, *Pseudotsuga menziesii*) that are considered for silvicultural climate change adaptation in managed forests. Foliage of mature trees was incubated at temperatures from 35-50 °C for up to 4 h to simulate daily heat maxima during heat waves. The maximum quantum yield (*F*_v_/*F*_m_) of photosystem II (PS II) of dark-adapted leaves was measured, because the PS II is particularly sensitive to heat and its functionality can decide on plant survival under heat.

*Fagus sylvatica* was much more tolerant to heat than *Pseudotsuga menziesii*, but weakly (albeit significantly) less tolerant than *Fagus orientalis*. Within its limits, *Pseudotsuga menziesii* showed high seasonal heat acclimation with constantly increasing tolerance during the growing season. *Fagus orientalis*, but practically not *Fagus sylvatica*, also acclimated to heat. This makes *Fagus orientalis* slightly superior over *Fagus sylvatica* in terms of heat tolerance, whereas the suitability of *Pseudotsuga menziesii* for silvicultural climate change adaptation is questionable. Strong heat acclimation, but also overall low heat tolerance, in *Pseudotsuga menziesii* might be the result of evergreenness, which requires the generation of both cold and heat tolerance during the year.

## 1. Introduction

There is multiple concern about the climate change tolerance of European beech (*Fagus sylvatica*), which responds to trends for increases in temperature and vapor pressure deficit (VPD) with reduced growth (Martinez del Castillo et al. 2022; Klesse et al. 2024). Compound heat and drought events, like during the multiyear-drought of 2018-2020, have resulted in increased mortality of *F. sylvatica* (Schuldt et al. 2020; Obladen et al. 2021; Rukh et al. 2023). Early leaf senescence is a typical response to hot droughts that is more widespread in *F. sylvatica* than in other temperate broadleaves of Europe (Frei et al. 2022) and can, at least partly, be connected to the high uncontrolled cuticular transpiration in this species (Van Gardingen & Grace 1992; Kerstiens 1995).

While much research has been dedicated to the influence of drought on the vitality of temperate tree species and on the occurrence of xylem hydraulic failure in particular (Wortemann et al. 2011; Schuldt et al. 2016; Herbette et al. 2021), heat was for a long time considered secondary for climate change-induced vitality reductions and increased tree mortality and has only recently been recognized as a critical factor that acts largely independently of drought stress (Konôpková et al. 2018; Kunert & Hajek 2022; Húdoková et al. 2022). While in many cases drought-tolerant tree species are at the same time also heat-tolerant, which makes sense from an evolutionary point of view as water shortages and high temperatures often coincide, systematic analysis of several temperate tree species showed that there are cases where tolerance to drought and heat are independent of each other (Hauck et al. 2025). For instance, drought tolerance (measured as *P*_88_, i.e. 88% loss of xylem conductivity due to embolism; Delzon & Cochard 2014) in temperate oak species decreases in the order *Quercus pubescens* > *Q. rubra* > *Q. robur*, while the heat tolerance of photosystem II (PS II) does not differ between these species (Hauck et al. 2025).

The heat sensitivity of plants is closely linked with PS II dysfunction and with reductions in CO_2_ assimilation in the Calvin cycle (Mohanty et al. 2012). CO_2_ assimilation is inhibited by heat, as Rubisco activase that is needed for the functioning of Rubisco (which is heat-stable in itself) is extremely heat-sensitive (Salvucci & Crafts-Brandner 2004). While this effect of heat on the Calvin cycle at weak to moderate heat even below 40 °C, is about reversible enzyme inactivation and not about structural damage (Haldimann & Feller 2005). Heat effects on PS II, however, are connected with structural damage and the repair mechanisms of PSII are particularly sensitive even to moderate heat (Mohanty et al. 2012). For the obvious reason that an intact PSII is essential for the light reaction of photosynthesis, the destruction of PS II in large parts of a tree’s foliage is equivalent to tree death due to the essential role of PS II for photosynthesis (Guadagno et al. 2017).

The mechanisms leading to PSII disintegration under heat are known in great detail (Allakhverdiev et al. 2008; Mohanty et al. 2012): The Mn-containing oxygen-evolving complex is a particularly heat-sensitive component of PSII (Nash et al. 1985; Enami et al. 1994). At temperatures >40 °C, the chlorophyll *a*-containing D1/D2 heterodimer that forms the core-protein of the PSII dissociates into two monomers (Lípová et a. 2010; Mohanty et al. 2012). Moreover, the light-harvesting complexes that contain chlorophyll *a* and *b* detach from the PSII core protein (Zhang et al. 2012). Heat also increases the fluidity and permeability of thylakoid membranes (Hüve et al. 2011) and induces structural changes in the thylakoids by grana destacking at temperatures around 35-45 °C (Gounaris et al. 1984; Georgieva 1999; Andersson et al. 2003). The generation of reactive oxygen species (ROS) onsets under moderate heat due to malfunction of the oxygen-evolving complex and intensifies PS II dysfunction, for example, by inhibiting the synthesis of the D1 protein (Hüve et al. 2011; Mohanty et al. 2012; Yamamoto 2016). The photosystem I (PS I) is much more heat-stable than the PS II, making the analysis of PS II chlorophyll fluorescence to an ideal tool for the study of heat tolerance in plants (Berry & Björkman 1980; Zhang et al. 2012).

Most of these mechanisms causing the heat sensitivity of PS II are obviously not directly related to drought sensitivity, which suggests that the examples of tree species where high or low heat and drought tolerance coincide (Hauck et al. 2025) are not the result of a common mechanism. However, there are also interdependencies between heat and drought tolerance, as transpirational cooling can prevent leaves from reaching critical temperatures (Marchin et al. 2023) and as the generation of cellular heat tolerance depends as in the case of drought tolerance on the availability of assimilates and can be favored by the presence of osmolytes (Papageorgiou & Murata 1995; Chen & Murata 2002).

Plants can acclimate to heat as a response to heat exposure in the field by the formation of heat shock proteins (HSP) that protect proteins from denaturation (Zhu et al. 2024) and by changes in membrane biochemistry. Both heat and cold acclimation include changes in the lipid composition of thylakoids and other cell membranes to maintain membrane fluidity and structural integrity (Velitchkova et al. 2009). At low temperatures, some lipids transition from the fluid phase to the solid gel phase earlier than others causing lateral phase separation and membrane dysfunction (Berry & Björkman 1980). Depending on lipid composition, temperatures of phase separation can widely vary (Pike & Berry 1980). A high share of unsaturated fatty acids can stabilize membrane integrity under heat (Zheng et al. 2011; Reszczyńska & Hanaka 2020). However, this is counteracted by ROS formation, which causes lipid peroxidation (Yamamoto et al. 2016). While lipid unsaturation is slow and requires high energy inputs and assimilates, changes in the hydrophilic head groups of glycerolipids constitute heat tolerance more quickly at lower energy demand (Zheng et al. 2011).

Heat acclimation of tree species during the growing season has hardly been addressed in the literature, so far (Posch et al. 2025). Húdoková et al. 2022 examined the PS II heat tolerance at two points in time in summer (in late June and early August) and found higher tolerance during the second measurement campaign in *Quercus petraea*. Petrik et al. (2023) conducted monthly measurements of PS II heat tolerance in *Picea abies* from May to September and found a significantly lower maximum quantum yield of PS II (*F*_v_/*F*_m_) in May compared to the following months after exposure to 48 °C for 30 min. Otherwise, series with several repeat measurements during the growing seasons have not yet been published. The studies of Kunert & Hajek (2022), Kunert et al. (2022), Münchinger et al. (2023), and Hauck et al. (2025) as well as others from other biomes (Tiwari et al. 2020; Slot et al. 2021) or from the North America part of the temperate forest biome (Endris & Rehm 2025) were selective measurements of PS II heat tolerance with the aim to compare different species in principle, but not at different points in time. Therefore, we also integrated the seasonal variability of PS II heat tolerance in order to capture potential variations due to heat acclimation during the growing season.

The limited drought tolerance of *F. sylvatica* (Leuschner 2020) has prompted considerations as to whether this species should be partly replaced by other putatively more drought-tolerant tree species as a means of the climate change adaptation of forest management (Bolte et al. 2009; Fuchs et al. 2024; Wrzesiński et al. 2024). Mostly exotic tree species are under debate as potential candidates for such adaptation approaches (Hauck 2023). Two species that have been suggested for the partial replacement of *F. sylvatica* are Oriental beech (*Fagus orientalis*) (Mellert & Šeho 2022) and Douglas fir (*Pseudotsuga menziesii*) (Isaac-Renton et al. 2014; Thomas et al. 2022).

The rationale of using *F. orientalis* for the climate change adaptation of Central European forests is the more southern distribution range compared to that of *F. sylvatica*, which suggests the tolerance to warmer and drier climate extremes (Mellert & Šeho 2022). *P. menziesii* is considered as a suitable candidate because of its origin from the temperate zone of western North America, where summers are naturally drier than in Central Europe (Paruelo et al. 1995; Orth et al. 2016). Recent studies, however, showed that both species have their limitations with regard to their suitability for climate change adaptation in Europe. Even though the drought tolerance of *P. menziesii* is higher compared to *Picea abies* (Vitali et al. 2017; Dietrich et al. 2025), there is growing evidence of drought-related growth reductions and mortality increases in *P. menziesii* in Europe (Sergent et al. 2014; Vejpustková & Čihák 2019; Cavelier et al. 2025) as well as in its native range in North America (Restaino et al. 2016; Leuschner & Meinzer 2024). Furthermore, comparative studies of the drought sensitivity of stemwood formation in introduced *F. orientalis* and native *F. sylvatica* at sites in Central Europe where both species co-occur revealed only minor differences between these species (Kohler et al. 2024). While *F. orientalis* was slightly superior in terms of summer drought tolerance, it was more susceptible to spring drought than *F. sylvatica* (Kohler et al. 2024).

In the present study, we analyze the heat tolerance of *F. sylvatica* in comparison with *F. orientalis* and *P. menziesii*. Methodologically, we build on Hauck et al. (2025), where PS II heat tolerance was tested at more realistic time spans of several hours of heat exposure, provoking damage at lower temperature than in previously published experiments on heat tolerance of *F. sylvatica* with heat treatments of 15-30 min (Kunert & Hajek 2022; Münchinger et al. 2023). After 4 h heat exposure, Hauck et al. (2025) showed a depression of *F*_v_/*F*_m_ at a leaf temperature of 45 °C in *F. sylvatica*, but a complete collapse of PS II (*F*_v_/*F*_m_ = 0) in *P. menziesii*. A similar result for *F. sylvatica* was obtained by Kurjak et al. (2019), who exposed leaves to rising heat in steps of 3 K for 30 min from 30 to 48 °C. Published data on the heat tolerance of *F. orientalis* are not available. While the study of Hauck et al. (2025) was limited to the comparison of different tree species that were mostly sampled only once during the growing season, we extended our analyses to a study of the seasonal variation in the heat sensitivity of *F*_v_/*F*_m_ in the present study.

Based on the similar drought tolerance of *F. sylvatica* and *F. orientalis* (Kohler et al. 2024) and on the fact that both species split relatively recently in evolutionary terms at the turn of the Pliocene to the Pleistocene (Jiang et al. 2022), we hypothesized (1) that the heat tolerance of *F. sylvatica* and *F. orientalis* does not differ significantly. Furthermore, we tested the hypotheses (2) that the heat tolerance of *P. menziesii* is consistently lower than that of either beech species and (3) that heat tolerance of all species increases over the growing season due to acclimation.

## 2. Materials and Methods

### 2.1. Plant material and climate in the collection area

Branches of mature forest trees were collected in the Black Forest and the Upper Rhine valley of southwestern Germany near Karlsruhe and Freiburg. *Fagus sylvatica* L. and *F. orientalis* Lipsky were sampled from a mixed stand (49°7’ N, 8°27’ E, 115 m a.s.l.) in the Hardtwald near Vorsenz ca. 9 km N of Karlsruhe (Kohler et al. 2024). The population of *F. orientalis* was founded in 1940 by planting trees of a provenance from the Greater Caucasus (Kurz et al. 2023). Branches of *Pseudotsuga menziesii* (Mirb.) Franco of unknown provenance were collected in the Black Forest near Simonswald (48°6’ N, 8°3’ E, 300 m a.s.l.) ca. 20 km NE of Freiburg and ca. 115 km S of Karlsruhe.

The study region in southwestern Germany is the warmest region of Germany. While temperatures in Freiburg and Karlsruhe are similar, Freiburg receives more precipitation due to its location directly west of the Black Forest. From 1990-2025, mean annual temperature amounted to 11.6 °C in Freiburg and 11.4 °C in Karlsruhe. Mean annual precipitation in this period was 890 mm in Freiburg and 760 mm in Karlsruhe. These data are based on the weather stations #1443 for Freiburg and #2522/#4177 (change of station location by the data provider in 2008) for Karlsruhe (German Meteorological Service, Climate Data Center). Annual heat maxima during 1990-2025 ranged from 32.4-40.2 °C in Freiburg and from 32.7-40.2 °C in Karlsruhe. These high temperature maxima in the study area suggest that the trees had opportunity for heat acclimation during the growing season. Samples were collected in 2025 in March, May, July, and October for *P. menziesii* and in April, May, July, August, and September for the *Fagus* species. Beech branches in April were collected soon after bud break when leaves were still delicately green, but had achieved their full size. The year of 2025 was no exceptional year in terms of mean and maximum temperatures in the study region (Fig. S1). With 11.6 °C in Freiburg and 11.7 °C in Karlsruhe, the mean temperatures in 2025 were very close to the average of 1990-2025.

To ensure that we captured seasonal acclimation without interferences by differences specific to tree individuals or the spatial variation in site conditions, we sampled always the same tree individuals and limited the study to one tree individual per species. We were well aware of the fact that this procedure could be questioned with respect to representativeness, but for *F. sylvatica* and *P. menziesii*, where published heat tolerance measurements with the same methodology were available from individual dates from different trees and sites (Hauck et al. 2025), the results were well comparable with the present observations. We collected three sun-exposed branches per tree and date at ca. 5 m height using telescope scissors. Branches were kept well-watered during transport to the laboratory and stored at 4 °C before use. We harvested leaves at random and mixed them from the three branches per species and incubated and measured a minimum of 6 leaves per combination of temperature and incubation time as replicates.

### 2.2. Heat treatment

We incubated complete leaves in a preheated water bath at 35, 40, 45, and 50 °C for 15, 30, 45, 60, 120, 180, and 240 min. We preferred this compared to using leaf disks to work as close to natural conditions as possible. Leaves were put in paper bags and sealed watertight in heat-stable plastic bags. This way, the leaves were sealed with some air and kept dry during incubation. For heating, an immersion boiling device (Sous Vide Stick Enfinigy, Zwilling, Solingen, Germany) was used. We checked that the temperature in the interior of the sample bags was the same as in the water bath with control measurements using temperature sensors (TC 319, Dostmann electronic, Wertheim, Germany) positioned in the sealed paper bags (Table S1). Fresh leaves were used for every combination of temperature and incubation time. In the case of *P. menziesii*, we used 1–3-year-old needles. We tested that the incubation procedure itself without the influence of heat did not affect *F*_v_/*F*_m_ by incubating leaves at 25 °C for 4 h (Table S2), as we consistently measured *F*_v_/*F*_m_ well above 0.8 (Murchie & Lawson 2013).

### 2.3. Chlorophyll fluorescence analysis

Leaves were kept in the dry paper bags at dark for at least 20 min after the heat treatment. Chlorophyll fluorescence was recorded using a Mini-PAM-II Photosynthesis Yield Analyzer (Walz, Effeltrich, Germany). The minimal level of chlorophyll fluorescence (*F*_0_) was measured using a weak measuring beam that was not sufficient to induce electron transport at PS II. Maximum fluorescence (*F*_m_) was recorded after applying a saturation pulse of 5000 µmol m^-2^ s^-1^ for 600 ms (Murchie & Lawson 2013). The maximum quantum yield of PS II (*F*_v_/*F*_m_) was calculated as *F*_v_/*F*_m_ = (*F*_m_ – *F*_0_)*/F*_m_.

### 2.4. Statistical analysis

Arithmetic means ± standard errors (SE) are presented throughout the paper. Statistical analyses were computed in R 4.4.1 software. Effects of tree species, temperature, and incubation time on *F*_v_/*F*_m_ were tested with generalized linear models (GLM) selecting a quasi-binominal distribution and a logit link function (Table 1). To test for the significance of seasonal heat acclimation, we focused on month and the interaction term of incubation time × month for the individual tree species in a separate GLM analysis (Table 2). Predictor selection was aided by checking for multicollinearity with the R package ‘performance’ 0.12.4. The variance inflation factors (VIF) varied between only 1.0 and 1.1 in the models used. In the GLM for *P. menziesii* at 45 and 50 °C in Table 2, we had to exclude the interaction term incubation time × month from the models, because they caused the VIF to increase to 6.7 and 21, respectively. Despite the exclusion of these terms and VIF of 1.0 for the predictors that remained in the GLM, these two models explained as much as 86% and 94%. Numeric parameters were rescaled before analysis in GLM by subtracted the sample mean from the individual values and dividing the result by the sample’s standard deviation using the ‘scale’ function in R.

**Table 1.**
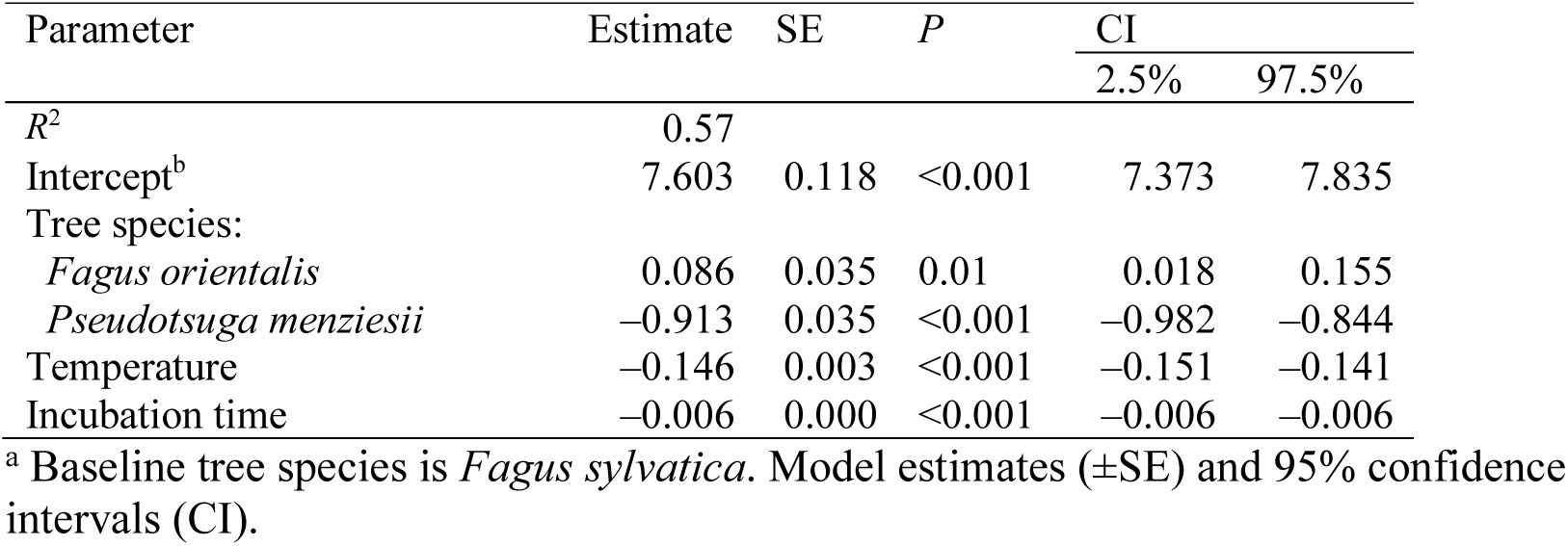
Generalized linear model for the effects of tree species, temperature and incubation time on the chlorophyll fluorescence yield of dark-adapted leaves^a^.

**Table 2.** Generalized linear models calculated separately for tree species and temperature for the effects of incubation time, month and the interaction term incubation time × month on the chlorophyll fluorescence yield of dark-adapted leaves^a^.

| Temperature | *F. sylvatica* | *F. orientalis* | *P. menziesii* |
| --- | --- | --- | --- |
| $R^2$ | | | |
| 35 °C | 0.30 | 0.53 | 0.67 |
| 40 °C | 0.23 | 0.30 | 0.75 |
| 45 °C | 0.41 | 0.62 | 0.94 |
| 50 °C | 0.47 | 0.47 | 0.86 |
| Intercept: |  |  |  |
| 35 °C | 1.582±0.009*** | 1.548±0.006*** | 1.452±0.015*** |
| 40 °C | 1.448±0.011*** | 1.451±0.013*** | 1.004±0.022*** |
| 45 °C | 0.694±0.040*** | 0.838±0.029*** | −2.637±0.119*** |
| 50 °C | 0.427±0.058*** | −0.200±0.055*** | 6.404±0.528** |
| Incubation time: |  |  |  |
| 35 °C | −0.012±0.009 | −0.009±0.006 | −0.109±0.015*** |
| 40 °C | −0.083±0.011*** | −0.122±0.013*** | −0.333±0.021*** |
| 45 °C | −0.436±0.004*** | −0.389±0.028*** | −3.750±0.160*** |
| 50 °C | −1.036±0.069*** | −0.904±0.062*** | −6.968±0.604*** |
| Month: |  |  |  |
| 35 °C | −0.098±0.009*** | −0.116±0.007*** | 0.314±0.015*** |
| 40 °C | 0.023±0.011* | 0.002±0.013 | 0.517±0.023*** |
| 45 °C | 0.323±0.039*** | 0.467±0.028*** | 0.380±0.046*** |
| 50 °C | 0.015±0.058 | 0.042±0.054 | 0.344±0.116** |
| Incubation time × Month: | | | |
| 35 °C | −0.002±0.009 | 0.001±0.007 | 0.036±0.015* |
| 40 °C | 0.065±0.011*** | 0.073±0.013*** | 0.019±0.022 |
| 45 °C | −0.042±0.039 | 0.075±0.028* | . |
| 50 °C | −0.082±0.068 | 0.095±0.061 | . |

<sup>a</sup> Estimates ± SE. Levels of significance: \* $P \leq 0.05$ , \*\* $P \leq 0.01$ , \*\*\* $P \leq 0.001$

## 3. Results

### 3.1. Influence of incubation time on heat tolerance

The duration of heat exposure, which was varied between 15 min and 4 h in our experiments, exerts a significant influence on PS II heat tolerance (Table 1). In the overall regression model for all data, incubation time had approximately half as much influence on reducing *F*_v_/*F*_m_ during the heat treatment as temperature (Table 1). The longer the heat treatment, the stronger was the response of *F*_v_/*F*_m_, as soon as a critical temperature threshold was crossed (Fig. 1). In the case of *P. menziesii*, a first slight reduction of *F*_v_/*F*_m_ started to occur at 35 °C and 2-4 h of heat exposure with treatment means of *F*_v_/*F*_m_ of 0.78-0.79 (or in other words by 5-6 %) compared to 0.83 in the control treatment at 25 °C and 4 h (Table S1). In *F. orientalis* and *F. sylvatica*, the first signs of decline in *F*_v_/*F*_m_ started at 40 °C and 3-4 h (0.77-0.79) compared to 0.82-0.83 in the control (25 °C, 4 h) (Table S1).

**Fig. 1.**
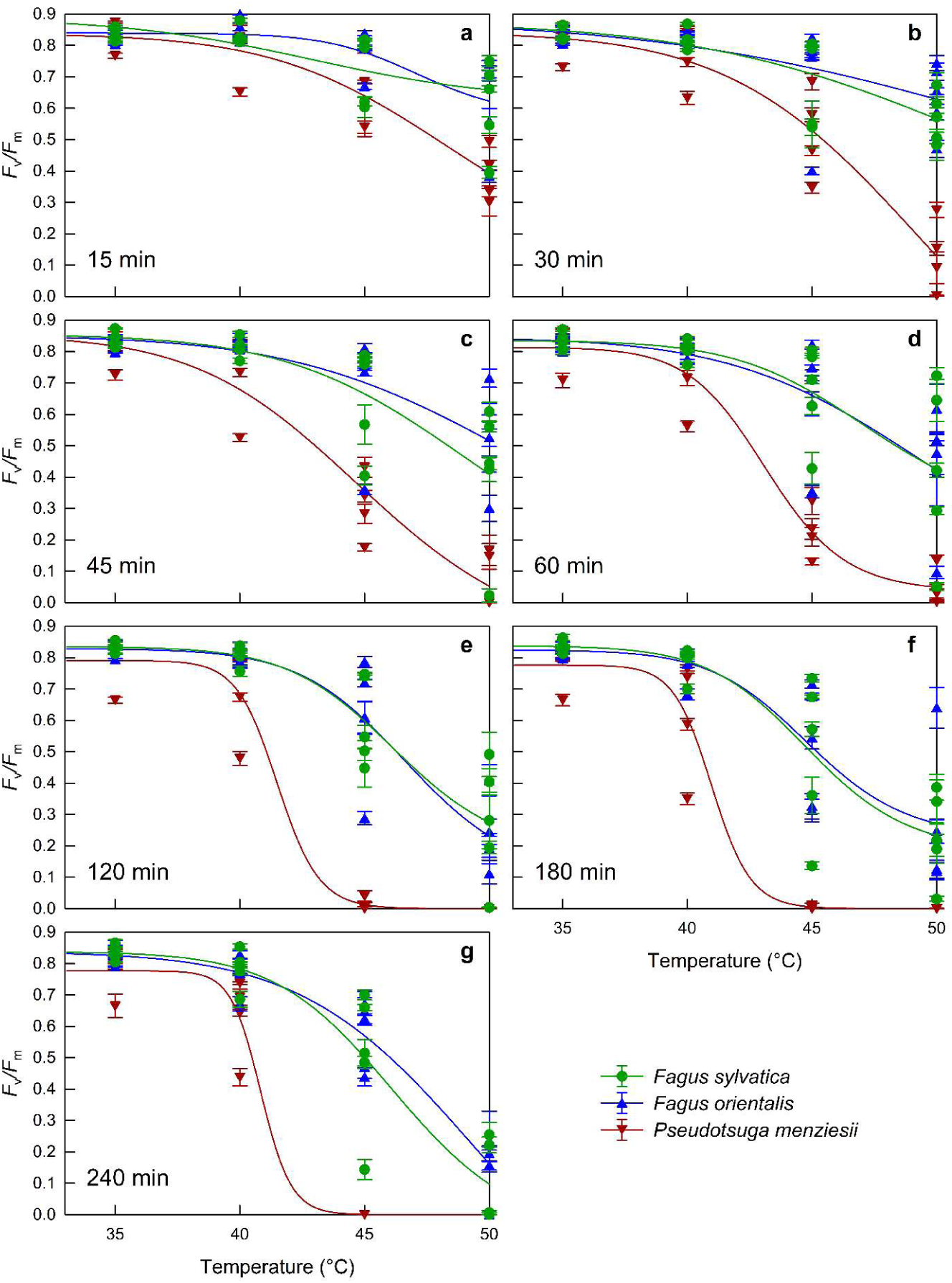
Maximum quantum yield of photosystem II (*F*_v_/*F*_m_) of leaves of *Fagus sylvatica*, *F. orientalis*, and *Pseudotsuga menziesii* incubated at 35 – 50 °C for up to 4 h. Differences between tree species are significant (see Table 1). Each marker (±SE) represents one combination of temperature and date.

In *P. menziesii*, the relationship of *F*_v_/*F*_m_ vs. temperature followed a clear logistic relationship at incubation times of ≥45 min, implying the occurrence of a steep decline of *F*_v_/*F*_m_ above a certain temperature threshold (Fig. 1). The slope gradient of this decline became steeper from 45 min via 1 h to 2 h of heat exposure, but remained constantly steep between 2 and 4 h (Fig. 1). In *F. orientalis* and *F. sylvatica*, the logistic shape of the regression line was less pronounced and occurred only during extended heat treatments of ≥2 h (Fig. 1).

### 3.2. Differences in heat tolerance between tree species

*F. orientalis* was slightly superior in terms of PS II heat tolerance over *F. sylvatica* (Fig. 1). Even though the difference in *F*_v_/*F*_m_ was small, it was statistically significant (Table 1). In *P. menziesii*, *F*_v_/*F*_m_ was much more strongly reduced by heat exposure than in the beech species (Fig. 1; Table 1). After 2 h of heat exposure, *F*_v_/*F*_m_ was close to 0 at 45 °C in *P. menziesii*, whereas it still amounted to 0.60±0.01 (mean of monthly mean values over all seasonal measurements) at this combination of temperature and incubation time both in *F. orientalis* and *F. sylvatica* (Fig. 1).

In separate regression analyses for the different tree species and temperatures (Table 2), the model intercepts for *F*_v_/*F*_m_ decreased much faster and more strongly in *P. menziesii* (from 0.81 at 35 °C to 0.18 at 50 °C) than in the two beech species (0.43-0.47). While the model intercepts were similar in *F. orientalis* and *F. sylvatica* at 35 °C (0.82) and 40 °C (0.81), they decreased faster in *F. sylvatica* than in *F. orientalis* at 45-50 °C (Table 2).

### 3.3. Heat acclimation

Our *F*_v_/*F*_m_ data showed clear signs of heat acclimation to different extents in the three studied tree species. The clearest indication of heat acclimation was found in *P. menziesii*. In most combinations of temperature and incubation time, *F*_v_/*F*_m_ increased in the different experiments during the growing season (Figs. 2c, 3c, 4c, 5c). In *P. menziesii*, this increase over the growing season included the control samples that were not exposed to any treatment in the laboratory at all. It was also observed in all heat-treated samples, expect those where the PS II collapsed, which happened in the treatments at 45 °C for ≥2h (Fig. 4c) and to 50 °C for ≥30 min (Fig. 5c).

**Fig. 2.**
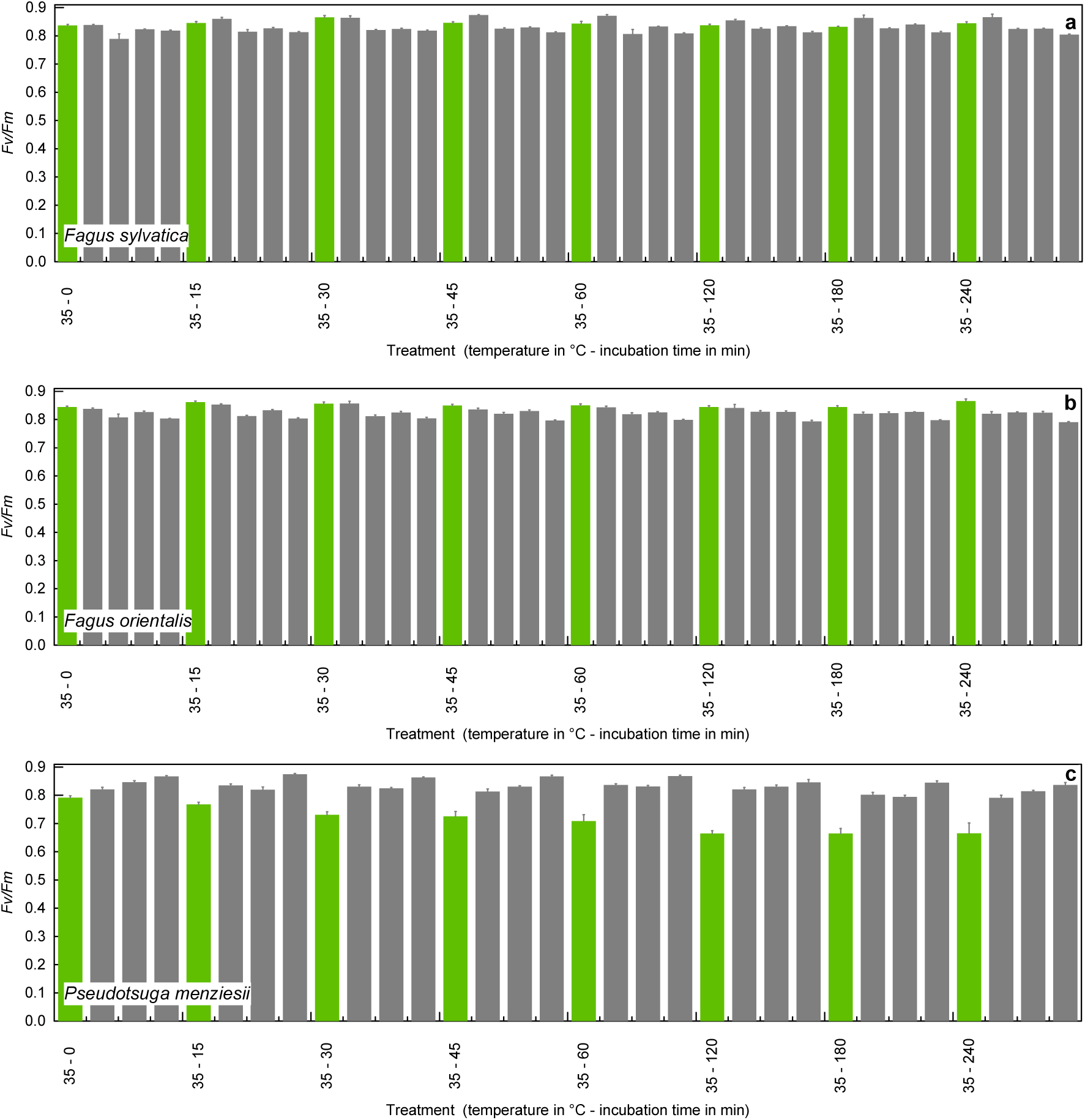
Seasonal variability of the maximum quantum yield of photosystem II (*F*_v_/*F*_m_) of leaves of (a) *Fagus sylvatica*, (b) *F. orientalis*, and (c) *Pseudotsuga menziesii* incubated at 35°C for up to 4 h. Columns represent measurements from April, May, July, August, and September in the deciduous *Fagus* species and from March, May, July, and October in the evergreen *P. menziesii*. Increases in *F*_v_/*F*_m_ over the growing season are statistically significant (see Table 2) The first column per temperature/incubation time combination is printed in green to improve visual clarity.

**Fig. 3.**
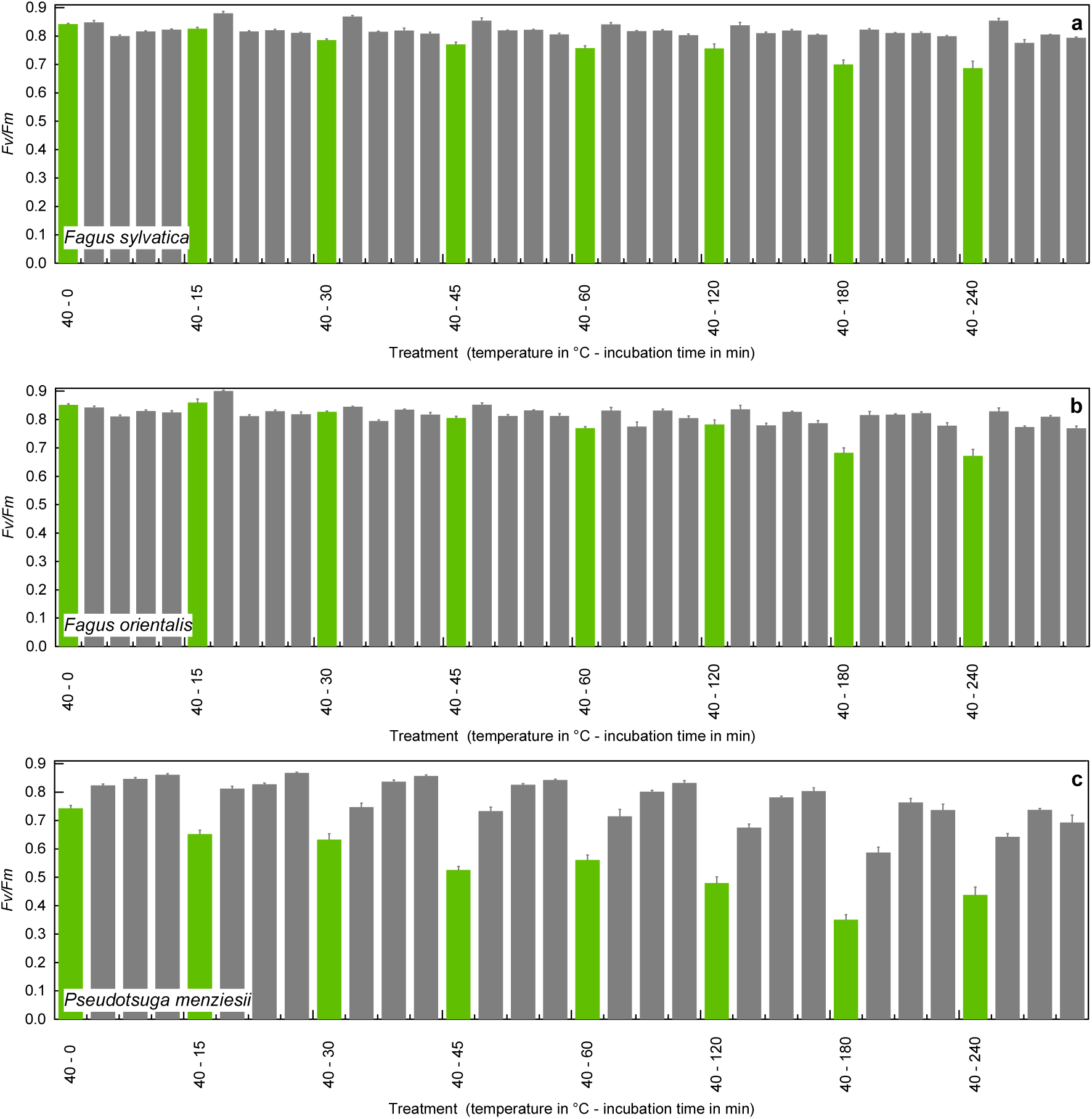
Seasonal variability of the maximum quantum yield of photosystem II (*F*_v_/*F*_m_) of leaves of (a) *Fagus sylvatica*, (b) *F. orientalis*, and (c) *Pseudotsuga menziesii* incubated at 40 °C for up to 4 h. Columns represent measurements from April, May, July, August, and September in the deciduous *Fagus* species and from March, May, July, and October in the evergreen *P. menziesii*. Increases in *F*_v_/*F*_m_ over the growing season are statistically significant (see Table 2).

**Fig. 4.**
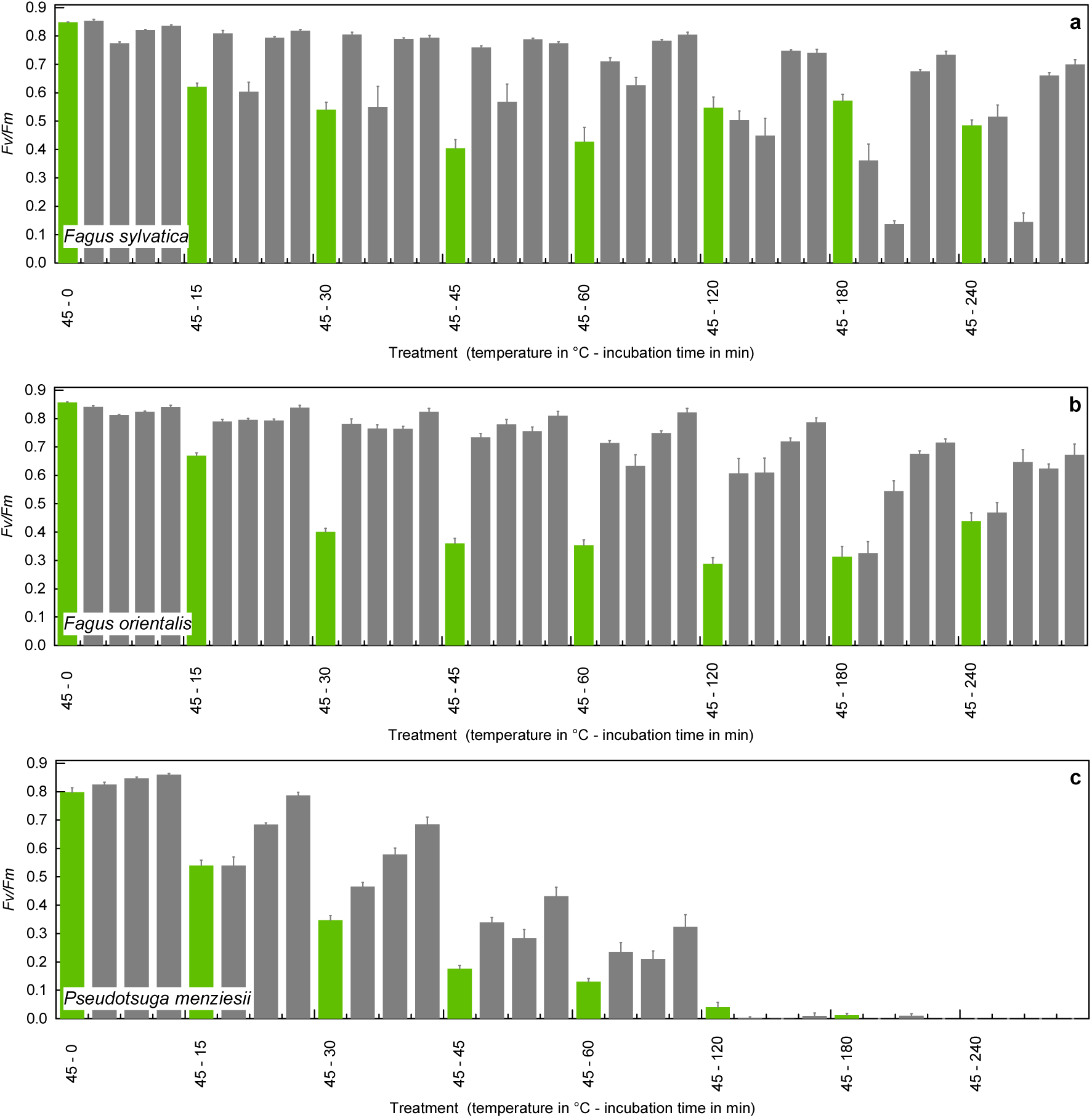
Seasonal variability of the maximum quantum yield of photosystem II (*F*_v_/*F*_m_) of leaves of (a) *Fagus sylvatica*, (b) *F. orientalis*, and (c) *Pseudotsuga menziesii* incubated at 45°C for up to 4 h. Columns represent measurements from April, May, July, August, and September in the deciduous *Fagus* species and from March, May, July, and October in the evergreen *P. menziesii*. Increases in *F*_v_/*F*_m_ over the growing season are statistically significant (see Table 2).

**Fig. 5.**
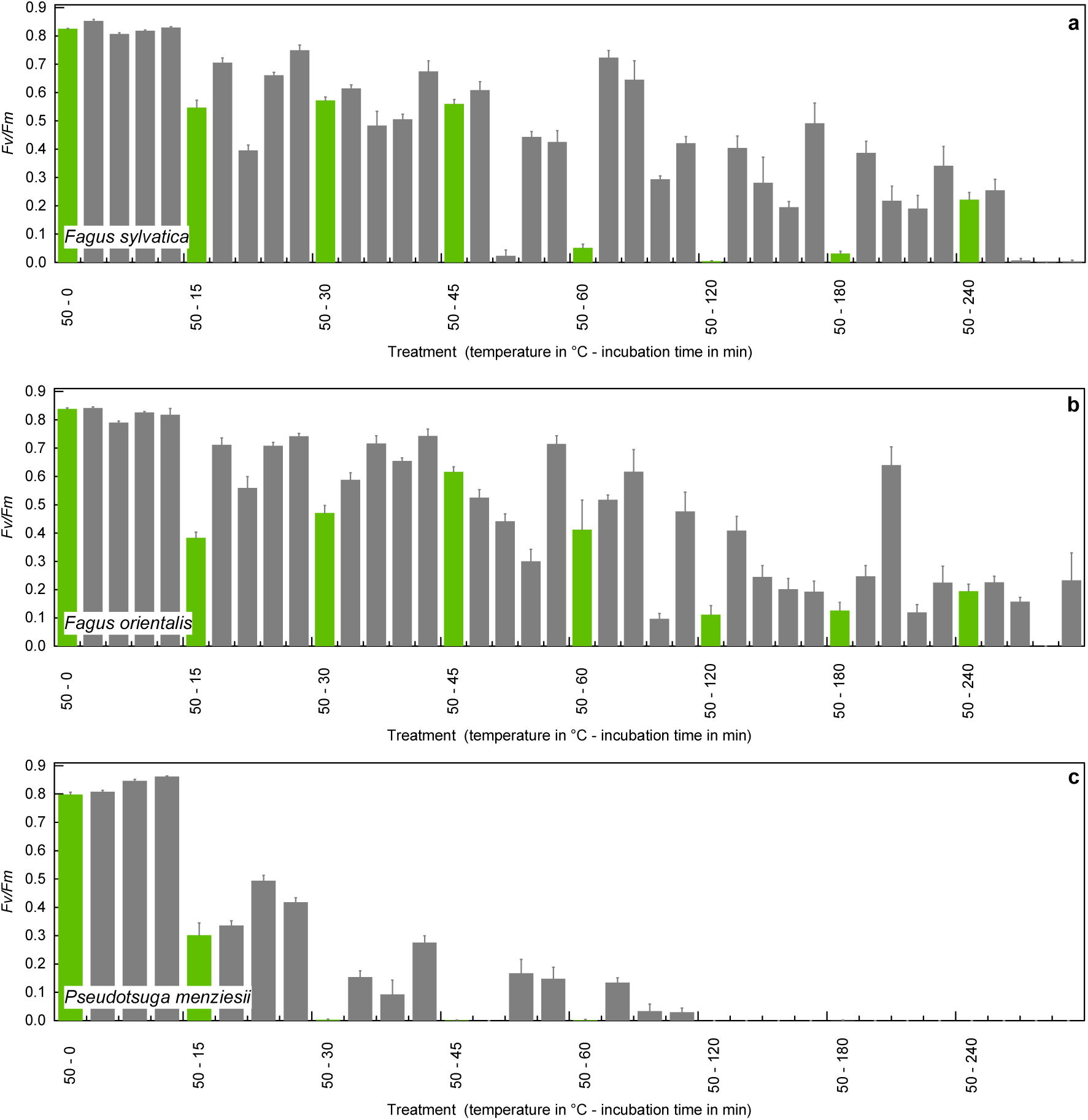
Seasonal variability of the maximum quantum yield of photosystem II (*F*_v_/*F*_m_) of leaves of (a) *Fagus sylvatica*, (b) *F. orientalis*, and (c) *Pseudotsuga menziesii* incubated at 50°C for up to 4 h. Columns represent measurements from April, May, July, August, and September in the deciduous *Fagus* species and from March, May, July, and October in the evergreen *P. menziesii*. Increases in *F*_v_/*F*_m_ over the growing season are statistically significant (see Table 2).

All these seasonal trends were significant in the regression models (Table 2). The increase in *F*_v_/*F*_m_ over the growing season was expressed by positive effects of the predictor ‘month’. Large differences in *F*_v_/*F*_m_ between the early and the late growing season are reflected by high model estimates. In combinations of temperature and incubation time with strong reductions of *F*_v_/*F*_m_ model estimates for ‘month’ decreased (Table 2).

The two beech species showed only marginal seasonal variability of *F*_v_/*F*_m_ at a temperature of 35 °C (Fig. 2a, b). If exposure to 40 °C exceeded 45 min, the newly formed, delicate green leaves in April showed declines of *F*_v_/*F*_m_ to 0.76-0.77 (45 min -2 h) and 0.69-0.70 (3-4 h) in *F. sylvatica*, whereas leaves of branches collected from May to September had *F*_v_/*F*_m_ values of 0.80-0.83, which was on the level of the control samples (Fig. 3a). The significance of this effect was corroborated by the positive effects of ‘month’ and ‘incubation time × month’ in the regression models (Table 2). In *F. orientalis*, such pattern with lower *F*_v_/*F*_m_ in April and stable *F*_v_/*F*_m_ during the remaining growing season was only observed in the 3 h and 4 h treatments at 40 °C (Fig. 3b), resulting in no significant effect of ‘month’, but a significant effect of the interaction term ‘incubation time × month’ (Table 2).

At 45 °C (all incubation times) and after short exposures up to 30 min at 50 °C, *F. orientalis* showed significant increases of *F*_v_/*F*_m_ during the growing season (Fig. 4b, 5b) similar to the widespread seasonal trends in *P. menziesii*. At 45 °C, this seasonal trend was significant in the regression model calculated for *F. orientalis* (Table 2). Incubation times ≥45 min at 50 °C caused a steady decline of *F*_v_/*F*_m_ from 0.52 at 45 min to 0.16 at 4 h (Fig. 5b). Under these conditions, a seasonal trend of *F*_v_/*F*_m_ failed to occur in *F. orientalis* (Fig. 5b; Table 2).

In contrast to *F. orientalis*, which was the most heat-tolerant study species, *F. sylvatica* showed a high variability of *F*_v_/*F*_m_ between the months at 45 and 50 °C (Fig. 4a, 5a). Despite a significant positive effect of ‘month’ in the regression model for *F. sylvatica* at 45 °C (Table 2), there were no steady increases in *F*_v_/*F*_m_ over growing season (Fig. 4a) like in the two other tree species.

## 4. Discussion

### 4.1. The role of incubation time for the validity of heat tolerance experiments

Our results confirm published findings of Hauck et al. (2025) that short-term experiments on the PS II heat tolerance of tree species of less than 1 h deliver less meaningful results compared to heat exposures over several hours. Yet, so far, heat treatments of 15 min (Krause et al. 2010; Tiwari et al. 2020; Slot et al. 2021; Kunert & Hajek 2022; Kunert et al. 2022; Endris & Rehm 2025), 30 min (Münchinger et al. 2023) or 40 min (Da Silva & Rossatto 2022, 2024) dominate the literature on the heat tolerance of tree species, published in seminal studies that led plant ecophysiologists on the way to broaden their perspective from tree hydraulics to the inclusion of heat in the investigation of climate change effects on trees.

Although even a test for 4 h remains a laboratory experiment under artificial conditions, it comes much closer to the duration of the warm noon and afternoon hours on a hot day than a short-term treatment. Therefore, and because neither in Hauck et al. (2025) nor in the present study, reductions in *F*_v_/*F*_m_ were caused in the control samples just by the 4 h incubation procedure without heat, our results from tests with up to 4 h are closer to ambient conditions and thus preferable over observations made in short-term experiments.

Our data (Fig. 1) suggest that even short-term experiments at 15 or 30 min are able to identify qualitative differences between species, but they are not appropriate to detect temperature thresholds for the thermostability of PS II under ambient conditions, as can be, for instance, taken from a comparison of the heat response of *P. menziesii* after 15 min (Fig. 1a) or 4 h (Fig. 1g) of heat exposure. Therefore, our results suggest that experiments on the heat tolerance of trees should generally be conducted over longer periods than has been customary in most studies to date. With our study, we confirm considerations of Neuner & Bucher (2023), who also emphasized the importance of longer incubation times in heat tolerance study and exposed plant leaves even over 8.5 h to heat.

### 4.2. Differences between tree species

In accordance with the second hypothesis, heat tolerance of *P. menziesii* was considerably lower than in the beech species. The complete collapse of the light reaction of photosynthesis in *P. menziesii* after 2 h exposure to 45 °C occurred at a temperature that exceeded the daily summer heat maxima in southwestern Germany measured in recent years by 5 K. However, it is well-known that canopy temperatures can exceed air temperatures by several K (Still et al. 2022). During heat waves in the native range of *P. menziesii* in western North America, air temperatures of 45 °C have already been recorded and even been exceeded (White et al. 2023; Leuschner & Meinzer 2024). In contrast to *P. menziesii*, *F*_v_/*F*_m_ in beech was only moderately reduced by heat exposure to 45 °C for 2 h, with 0.60±0.09 in *F. orientalis* and 0.60±0.06 in *F. sylvatica*. It is well documented in the literature that such moderate declines in *F*_v_/*F*_m_ can be overcome by repair mechanisms and can thus often be reversed (Kreslavski et al. 2008), whereas complete PS II collapse indicated by *F*_v_/*F*_m_ at or close to 0.00 is irreversible (Dascaliuc et al. 2007). The low heat tolerance of *P. menziesii* is in line with published values from heat treatments of up to 4 h in needles of mature *P. menziesii* from one sampling date (Hauck et al. 2025) and with high heat sensitivity of CO_2_ gas exchange in 4-year old *P. menziesii* (Duarte et al. 2016). The two beech species reached *F*_v_/*F*_m_ values where recovery is extremely unlikely not below 50 °C.

In contrast our first hypothesis, PS II heat tolerance in *F. orientalis* was significantly higher than in *F. sylvatica*. Though the difference between these species was minor, it could contribute to higher probability for survival during heat waves. The small, but significant superiority of *F. orientalis* in heat tolerance, matches with a trend towards higher mean annual temperatures and maximum temperatures in the distribution area of *F. orientalis* compared to that of *F. sylvatica* (Mellert & Šeho 2022).

### 4.3. Heat acclimation

*F. orientalis* also showed higher acclimation capacity to heat over the growing season than *F. sylvatica*. The seasonal heat acclimation in *F. orientalis* was most effective in the critical temperature ranges of 45 °C and, if exposed briefly, of 50 °C, before severe, putatively irreversible reductions in *F*_v_/*F*_m_ occurred. It is plausible to assume that the higher capacity for heat acclimation in *F. orientalis* is an important factor to constitute heat tolerance in this species. The significantly lower capacity for seasonal heat acclimation in *F. sylvatica* is in line with its more northern distribution range and the split of these two *Fagus* species in a cooling phase at the turn of the Pliocene to the Pleistocene (Jiang et al. 2022). Under the present rapid climate change in Central Europe, the lower capacity of *F. sylvatica* for heat acclimation over the growing season (that only partly confirms the third hypothesis) is clearly a disadvantage in comparison with *F. orientalis*. Especially *F. sylvatica* and to a lower extent *F. orientalis* would be vulnerable to extreme heat waves of right after bud break during the maturing process of the newly unfolded leaves. Such a temperature scenario is not realistic at present, but might become a problem in future, if unfavorable high-emission scenarios for greenhouse gas emissions become effective.

Despite its overall low heat tolerance, *P. menziesii* steadily increased its heat tolerance throughout the growing season. This significant seasonal heat acclimation is most likely related to the fact that, like other temperate and boreal conifers, *P. menziesii* undergoes intensive cold acclimation in winter (Chang et al. 2021), so that the heat tolerance in the evergreen needles has to be rebuilt every year. Energy and assimilate costs for temperature acclimation increase with the amplitude of temperature that has to be covered (Zheng et al. 2011). This means that over the course of the year, *P. menziesii* has to invest in changes in the lipid composition of the cell membranes both for cold and heat acclimation as well as in other measures to generate heat tolerance, like HPS production (Yoshida et al. 2011). The need to acclimate to both winter and summer conditions might also be crucial for the overall limited heat tolerance of *P. menziesii*, but also of other evergreen conifers, including *Abies alba*, *Picea abies*, and *Pinus sylvestris* (Kunert et al. 2022; Hauck et al. 2025). *F. orientalis* and *F. sylvatica* as deciduous broadleaves start into the growing season with newly formed leaves without a legacy of lipids for cold acclimation from winter.

While we aimed at analyzing heat tolerance over the growing season, Posch et al. (2025) pointed out that plants can acclimate at different temporal scales, a topic which is as yet largely unexplored in temperate tree species. However, heat acclimation of *F*_v_/*F*_m_ has been found in wheat (Posch et al. 2022) and in the tropical broadleaved Araliaceae tree *Polyscias elegans* (Zhu et al. 2024) within hours. It is obvious that these short-terms responses to heat must base on different mechanisms than the seasonal acclimation, which we primarily attribute to changes in lipid composition of the cell membranes (Velitchkova et al. 2009).

### 4.4. Management implications

The significantly lower heat tolerance of *P. menziesii* than of *F. sylvatica*, which is combined with a moderate drought tolerance (Leuschner & Meinzer 2024) does not support the use of *P. menziesii* for the climate change adaptation of managed forests in Central Europe. *P. menziesii* is not a species that can increase the tolerance or resilience of broadleaved temperate forests with a near-natural tree species composition (e.g. with *Fagus*, *Quercus*, and *Acer*; Hauck et al. 2025) in Central Europe to hot droughts. Concerns about the suitability of *P. menziesii* for the climate change adaptation of European forests have been uttered earlier (Hauck 2023; Cavelier et al. 2025; Leuschner 2025). The decision to grow *P. menziesii* is a mere economic one. Against the background of its ecophysiology, *P. menziesii* is also not recommended in terms of risk diversification, if compared to broadleaved native European forest communities. Growing *P. menziesii* in moist mountain forests, as has been suggested recently in the light of the species’ limited drought and heat tolerance (Leuschner 2025), can also only be justified as an economic decision and should not been seen as a means of climate change adaptation, because at such sites *F. sylvatica* can even benefit from warming at sufficient moisture supply (Dulamsuren et al. 2017).

For *F. orientalis*, the superiority both in terms of heat tolerance (this study) and drought tolerance (Kohler et al. 2024) is statistically significant, but minor. It questionable if these minor ecophysiological advantages under a warmer and drier climate outweigh the risks for biodiversity due to the introduction of this tree species outside its native range. In the case of the introduction of *F. orientalis* to Central Europe, these consequences also concern beech itself and not only dependent biota, as *F. orientalis* and *F. sylvatica* readily hybridize and thus the widespread introduction of *F. orientalis* to Central Europe would cause irreversible change to the genome of *F. sylvatica* (Budde et al. 2023; Kurz et al. 2023; Hrivnak et al. 2024). Therefore, more detailed knowledge of the tolerance of *F. sylvatica* and potential replacement tree species to climate change-induced weather extremes is needed (Huth et al. 2025).

## 5. Conclusions

Our study suggests that seasonal heat acclimation is a key feature for the mechanistic understanding of the thermotolerance of different tree species. The need to acclimate both to cold and heat in evergreen species by recurrent changes in the chemical composition of the cell membranes could explain the generally lower heat tolerance of *P. menziesii* as an evergreen conifer. The study of cumulative effects of repeated heat waves and of recovery for heat damage could be next steps in the research on the heat tolerance of temperate tree species.

## Author contributions

MH and CD developed the concept and wrote the paper. GS, KK, OS, YL, LP, and LS carried out field and laboratory work. MH analyzed the data.

